# Attractor neural networks with double well synapses

**DOI:** 10.1101/2023.07.17.549266

**Authors:** Yu Feng, Nicolas Brunel

## Abstract

It is widely believed that memory storage depends on activity-dependent synaptic modifications. Classical studies of learning and memory in neural networks describe synaptic efficacy either as continuous [1, 2] or discrete [2–4]. However, recent results suggest an intermediate scenario in which synaptic efficacy can be described by a continuous variable, but whose distribution is peaked around a small set of discrete values [5, 6]. Motivated by these results, we explored a model in which each synapse is described by a continuous variable that evolves in a potential with multiple minima. External inputs to the network can switch synapses from one potential well to another. Our analytical and numerical results show that this model can interpolate between models with discrete synapses which correspond to the deep potential limit [7], and models in which synapses evolve in a single quadratic potential [8]. We find that the storage capacity of the network with double-well synapses exhibits a power law dependence on the network size, rather than the logarithmic dependence observed in models with single well synapses [9]. In addition, synapses with deeper potential wells lead to more robust information storage in the presence of noise. When memories are sparsely encoded, the scaling of the capacity with network size is similar to previously studied network models in the sparse coding limit [2, 10–13].

## 1 Introduction

Synapses can change their strength in response to neuronal activity [14–18], and synaptic changes are thought to be critical for the brain to build memories [19]. A popular theoretical framework for studying learning and memory is the attractor neural network scenario [20–22]. In this class of models, memories correspond to the stable fixed points of the neuronal activity dynamics. External stimuli can modify the synaptic connectivity of the network following certain plasticity rules, and these synaptic modifications can create new stable fixed points of the neuronal dynamics. An external input is said to be stored in the network if there exists a stable fixed point that is highly correlated with it. An extensively studied question is the storage capacity of such networks [23–25], i.e. the number of stored memories, or the amount of information that can be stored. One key question in theoretical neuroscience is to find biologically plausible learning rules that lead to reasonably large storage capacities.

Multiple synaptic plasticity experiments have suggested that the strength of individual synapses is modified in a discrete manner. A few experiments have shown that synaptic plasticity is well described by switches between two discrete states (all-or-none), rather than by arbitrary continuous changes in efficacy [26, 27]. In addition, super-resolution imaging experiment have shown that dendritic spines contain a small discrete (1-4) number of nanomodules (i.e., clusters of receptors) [5], consistent with plasticity experiments and indicating that synapses are quantized. This picture of discrete synapses is in contrast with most studies of learning and memory in neuronal network models, in which synapses are taken to be real continuous variables.

Multiple efforts have been made to compute the storage capacity of networks with discrete synapses. Sompolinsky studied a network with a specific binarized Hebbian rule [28], and showed its capacity is close to the capacity of the Hopfield network [1]. Sompolinsky’s model assumes synapses still can store all continuous changes in synaptic efficacy during the learning phase, and synapses get clipped after they have learned all patterns. In contrast, Tsodyks [29], and Amit and Fusi [7, 30] introduced online learning rules in which synapses are always discrete. These studies found that discreteness of synapses can cause a large drop in storage capacity [29, 30], unless stored patterns become extremely sparse [7, 31]. Later studies introduced models with hidden states [32–35], but it remains unknown how these synaptic plasticity models perform in attractor neural networks.

While theoretical work has focused on either fully analog or fully discrete synapses, recent experimental results suggest an intermediate scenario in which the synaptic efficacy can be described by a continuous variable, but whose distribution is peaked around a small set of discrete values [6]. Motivated by this data, we introduce and explore a model in which synapses are described by a continuous variable that evolves in a potential with multiple minima (‘potential wells’, see Fig. 1A for an example with two wells). Synapses remain in the same potential well when receiving weak or short-lasting stimuli, but can switch to a new state when receiving strong or long-lasting stimuli. Our analytical and numerical results show this model can connect models with discrete synapses which correspond to the deep potential limit [7], and models in which synapses evolve in a single quadratic potential [8]. We show that the storage capacity of the network with double well synapses (and thus a bimodal weight distribution) has a power law dependence on network size. In contrast, models with a single-well potential show a logarithmic dependence when the parameters of the potential are not allowed to vary with network size *N*, as shown by previous studies [9]. we also show that the networks with double well synapses store information more robustly in the presence of noise.

**Figure 1.**
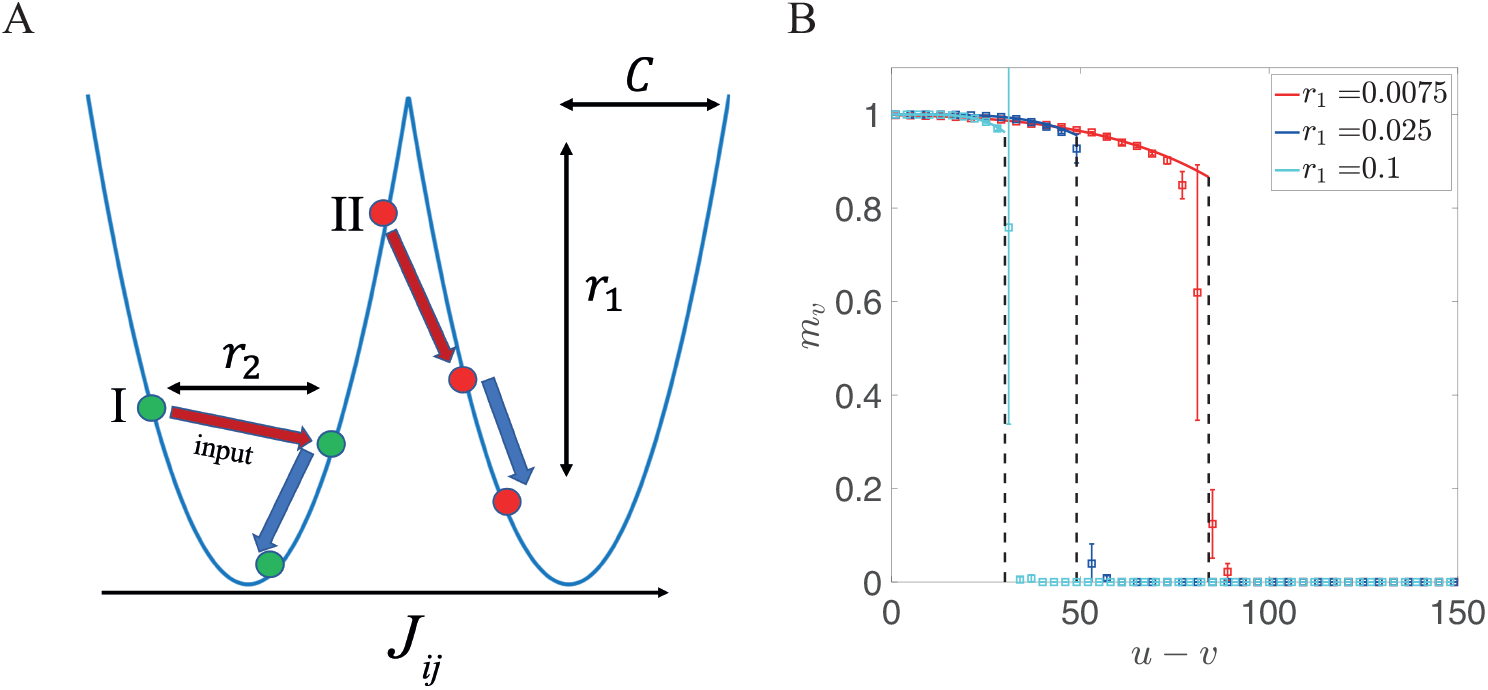
(A) Sketch of the synaptic model. In the presence of an external input, a synapse can stay in the same well (I), or jump into another well (II), depending on its current state and the amplitude of the input. *C* describes the distance between the two wells, while *r*_1_ characterizes its depth. The larger *r*_1_, the faster synapses decay towards the minima of the potential. (B) Overlap input presented at time *v* and its corresponding attractor state, *m*_*v*_, as a function of time elapsed since presentation of this input *u* − *v*, for different values of *r*_1_. The solid lines represent the theoretical prediction and squares represent the simulation results with a network of size *N* = 30, 000 (mean and standard deviation computed over ten independent realizations). Dashed lines mark the storage capacity where *m*_*v*_ vanishes. For all lines, the value of *C* is chosen to optimize storage capacity. Other parameters are: *r*_2_ = 1, *r*_3_ = 0, *θ* = 0, *f* = 0.5, *c* = 0.05.

## 2 Model and methods

We study a sparsely connected network of *N* binary neurons whose states *V*_*i*_ = 0, 1 (*i* = 1, …, *N*) are given by a parallel update rule:

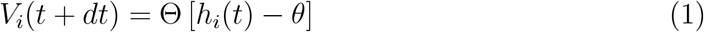

where *dt* ≪ 1 is the timescale of neuronal dynamics relative to synaptic dynamics, *θ* is a threshold, Θ is the Heaviside step function, and *h*_*i*_(*t*) is the local field of neuron *i*:

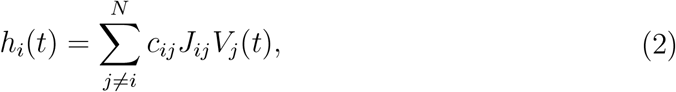

where *J*_*ij*_ is the strengths of the synapse from neuron *j* to neuron *i*, and *c*_*ij*_s are i.i.d Bernoulli random variables, *c*_*ij*_ = 1 with probability *c* ≪ 1, and 0 otherwise.

Synapses are assumed to be evolving on a much slower time than neuronal dynamics, and synaptic dynamics are supposed to be driven by external inputs that drive the network to specific states. Each synapse evolves according to

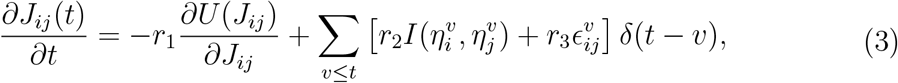

where *U* is a potential function with multiple minima, 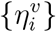 are i.i.d random binary variables that represent the state of the neuron imposed by external inputs at time *v* ∈ ℤ, with 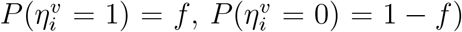, and 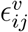 are i.i.d random Gaussian variables. Note that the chosen time unit is the interval between presentations of two successive external inputs.

In Equation (3), the dynamics of *J*_*ij*_ is determined by three terms, of respective strengths *r*_1_, *r*_2_ and *r*_3_. The first term determines the dynamics of the synapse in the absence of inputs. For simplicity we use a double well potential *U* in which each well is given by a quadratic function,

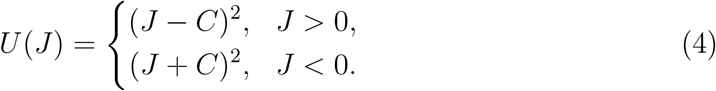

Thus, in the absence of inputs, each synapse decays to one of the two potential wells - a low efficacy state *J*_*ij*_ = −*C*, or a high efficacy state *J*_*ij*_ = *C* - exponentially, with a time constant 1*/*(2*r*_1_). Small values of *r*_1_ describe shallow potentials (and hence long decay times towards the minima) while large values of *r*_1_ describe deep potentials (and hence fast decay times towards the minima).

The second term 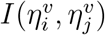 in Equation (3) is activity-dependent. For binary neurons, the function *I* can take at most 4 values, one for each combination of pre and post-synaptic activity. We study two scenarios: In the **balanced input** case, the coding level is *f* = 0.5, and we use

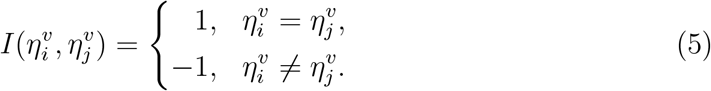

Note that for *U* = 0 (i.e. the flat potential case), and in the absence of noise, the model leads to a connectivity matrix that is identical to the Hopfield model [1]. This model however suffers from a ‘blackout’ catastrophe after a critical number of inputs, all memories suddenly becomes irretrievable. In the case of a potential with a single well (*C* = 0) the model leads to a connectivity matrix that is identical to a ‘palimpsest’ model introduced by Mézard et al [8]. This model instead can retrieve the most recently presented memories, but forgets memories some time after they have been presented.

In the **sparse input** case, the coding level is *f* ≪ 1. In this case it is no longer a good idea to use a balanced function *I*, since this choice would lead to the vast majority of synapses being in the low efficacy potential well. In this scenario, we use two different *I* functions. The first one is inspired by the Tsodyks-Feigelman model [10, 36]

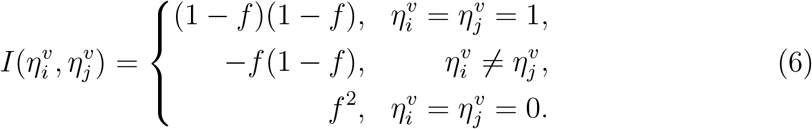

In the sparse input case, we also use another *I* function, motivated by the plasticity rule in [7]:

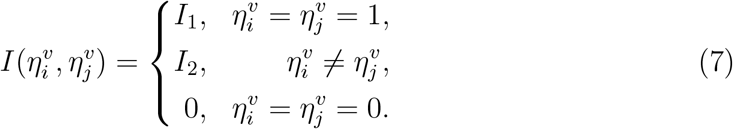

where *I*_1_ *>* 0 sets the strength of potentiation when both pre and post-synaptic neurons are active, *I*_2_ *<* 0 sets the strength of depression when one neuron is active while the other is inactive, and non changes occur when both neurons are inactive. Here, we focus on *I*_1_ = 2*C, I*_2_ = − *f* (1 − *f*). The rationale is that when the distribution of synaptic strengths is peaked around ± *C*, a potentiation strength of 2*C* leads to a transition from a lower to a higher state with high probability. When the connection is depressed, *I*_2_ takes the same value as in Equation (6), leading to a small transition probability from a higher to a lower state. This scheme mirrors the learning rule described in [7] for binary synapses.

The third term in Equation (3) is the noise term caused by fluctuations. These fluctuations could be of multiple origins: Fluctuations of neural activity; or fluctuations in the state of the synapse, due to fluctuations in vesicle release, numbers of activated receptors at the post-synaptic side, and fluctuations in biochemical reactions involved in synaptic efficacy changes.

Once the input patterns 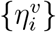 are given, the learning dynamics (3) is determined by four non-negative parameters: *C* determines the distance between two minima of *U* ; *r*_1_ how strongly the potential *U* affects the dynamics (i.e. how fast synapses decay to the minima of the potential); *r*_2_ is the input signal strength and *r*_3_ is the noise strength. In the presence of inputs, a synapse can stay in its current state, or jump into another state depending on its position in the potential well. This dynamics is illustrated in Figure 1A.

With such a synaptic plasticity dynamics and appropriate parameters, the network described by Equation (1) have multiple fixed points, that are close to the patterns imposed by the external inputs that were presented most recently. A pattern is said to be stored in memory if a network configuration close to the pattern is a stable fixed point of the dynamics. For this online synaptic plasticity rule eq. (3), we found that the most recently shown patterns *η*^*u*^, *η*^*u*−1^, …, *η*^*u*−*p*^ can be retrieved as stable fixed point attractors at a given time *u* ∈ ℤ, while older patterns cannot be retrieved. *p* is defined as the maximal number of patterns that are still retrievable or, equivalently the maximal age at which patterns can still be retrieved. This scenario is similar to previously studied attractor neural network models with online learning and decay term in the synaptic updates [8, 37], or networks with binary synapses [29, 38, 39] in which memories are forgotten exponentially.

In order to calculate the storage capacity for the network defined by eqs. (1) and (3), we consider a network at a particular time *u* when pattern *u* has just been presented, and investigate whether a pattern shown in the past *v < u* is still stored in the connectivity matrix. To determine whether there still exists an attractor state close to pattern *v*, we introduce as an order parameter the overlap between the pattern shown at time *v* and network state at time *u*:

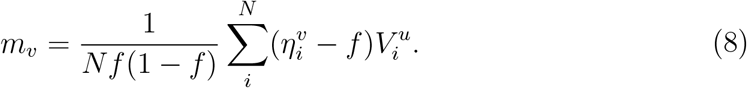

The order parameter *m*_*v*_ measures the retrieval quality of the pattern *η*^*v*^ stored in memory. States with *m*_*v*_ ∼ 1 mean that the network can correctly retrieve the stored memory. In practice, in numerical simulations, the network is initialized at 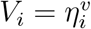, and the dynamics is run until convergence to a fixed point. Simulations reveal that sometimes, the network converges to the most recently shown pattern instead of the pattern *v*. Thus, we need to consider also the overlap with the most recently shown pattern, *m*_*u*_. In the balanced input case, we use analytical calculations to compute the mean-field equations for the overlap *m*_*v*_, and whether solutions to these equations with *m*_*v*_ *>* 0 are stable with respect to perturbations in the direction of the most recently shown pattern [37]. Details of analytical calculations are described in Appendix I and II. These calculations relie on approximating distributions of synaptic inputs, conditioned on 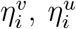, by Gaussian distributions. In the sparse input case, we used numerical simulations only, since this approximation is not accurate in this case for the values of *f* and *N* used in this paper.

The two order parameters *m*_*u*_ and *m*_*v*_ can be obtained by calculating the distribution of the local field *h*_*i*_ at the time *t* = *u*. In the large *N* limit (i.e., *cN* ≫ 1), the local field defined in eq. (2) follows a Gaussian distribution and its mean and variance are determined by the distribution of *J*_*ij*_. With the synaptic dynamics given by eq. (3), the probability density function of *J*_*ij*_ is determined by eqs. (16) and (17). The time-dependent distribution of *J*_*ij*_ can be solved numerically, and its mean value and standard deviation are given in eqs. (25) and (26). Then, the local field *h*_*i*_ can be written as a function of order parameters *m*_*v*_, *m*_*u*_ as shown in eqs. (33) and (34).

As shown in Appendix, *m*_*v*_, *m*_*u*_ are determined by two self-consistent equations:

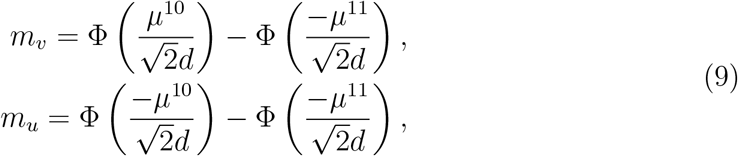

where 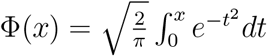, and *μ*^11^, *μ*^10^, *d* are functions of *m, m* defined in eqs. (33) and (34) (see Appendix II for details). Thus, *m*_*v*_ and *m*_*u*_ can be obtained solving numerically eqs. (16) to (19), (25), (26), (33) and (34). In practice, one can fix *v* and gradually increase *u* until there is no solution satisfying *m*_*v*_ ≠ 0, or this solution becomes unstable when a small overlap *m*_*u*_ is present. The storage capacity *p* is then obtained at the maximum value of *u* − *v* where a stable solution *m*_*v*_ ≠ 0 exists.

## 3 Results

### 3.1 Learning and retrieval dynamics for balanced input in the absence of noise

We start by considering the balanced input case (*f* = 0.5), and zero noise (*r*_3_ = 0). Without loss of generality, we can fix the value of *r*_2_ = 1, since the dynamics are invariant with respect to the transformation

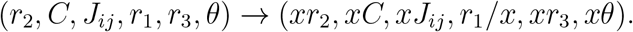

We also set *θ* = 0, since in this case the distribution of synaptic weights is symmetric around zero. For a given set of parameters, we use both mean field and numerical simulations to compute the overlap of a retrieved memory as a function of its age, and the storage capacity, defined as the maximum age at which a memory can be retrieved. Figure 1B shows the overlap of a memory learned at time *v, m*_*v*_ as a function of age (time *u* − *v* elapsed since the pattern was learned), for different potential depths *r*_1_ (other parameter values are indicated in the caption). We can see that the overlap decreases with age until it drops abruptly. The figure shows that analytic results are in good agreement with simulations, using a network with 30,000 neurons. It also shows that increasing *r*_1_ (i.e. the depth of the potential wells) decreases storage capacity. This is consistent with previous observations that clipping synaptic strength reduces capacity [3], and that online plasticity rules with discrete synapses typically have much lower capacity than rules with continuous synapses [7, 29, 30].

The storage capacity *p* thus strongly depends on the depth of potential wells as measured by *r*_1_, but it also depends on the width of the potential wells *C* relative to the jumps caused by input patterns. To investigate the storage capacity of the network, we begin by examining the single potential case, where the width of the potential wells *C* is set to zero.

#### Single Well Potential

When *C* = 0, the potential defined in Equation (4) becomes a single parabolic function. In this case, each synapse decays to zero exponentially with a time constant *τ* = 1*/*(2*r*_1_) in the absence of inputs, according to the dynamics defined in Equation (3). This scenario is similar to the previously investigated palimpsest model [8, 40] or the pure forgetting model of ref. [9]. The storage capacity *p* in this case depends on the time constant *τ* as [9]:

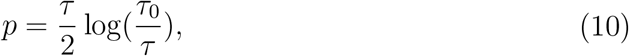

where *τ*_0_ = 2*A*(*f*)*N*, and where *A*(*f*) is a function of the coding level and thus is a constant in the balanced input case. When *τ > τ*_0_, no patterns are retrievable, similar to the Hopfield model scenario when the number of patterns exceeds the critical value. When *τ < τ*_0_, the model can always retrieve recent patterns and becomes a palimpsest model. By allowing the time constant *τ* in Equation (10) to scale linearly with the network size *N*, the storage capacity *p* can have a linear dependence with *N*, as investigated in previous studies [8, 40]. However, this scaling assumes an unrealistically large synaptic decay time for large networks. In this study, we consider the decay constant *τ* or potential depth *r*_1_ to be a property of individual synapses, rather than being dependent on network size, and thus *r*_1_ ∼ *O*(1) in the large *N* limit. In this scenario, the weight dynamics with a single well potential only leads to a storage capacity that increases logarithmically with *N*, as shown in Figure 2C.

**Figure 2.**
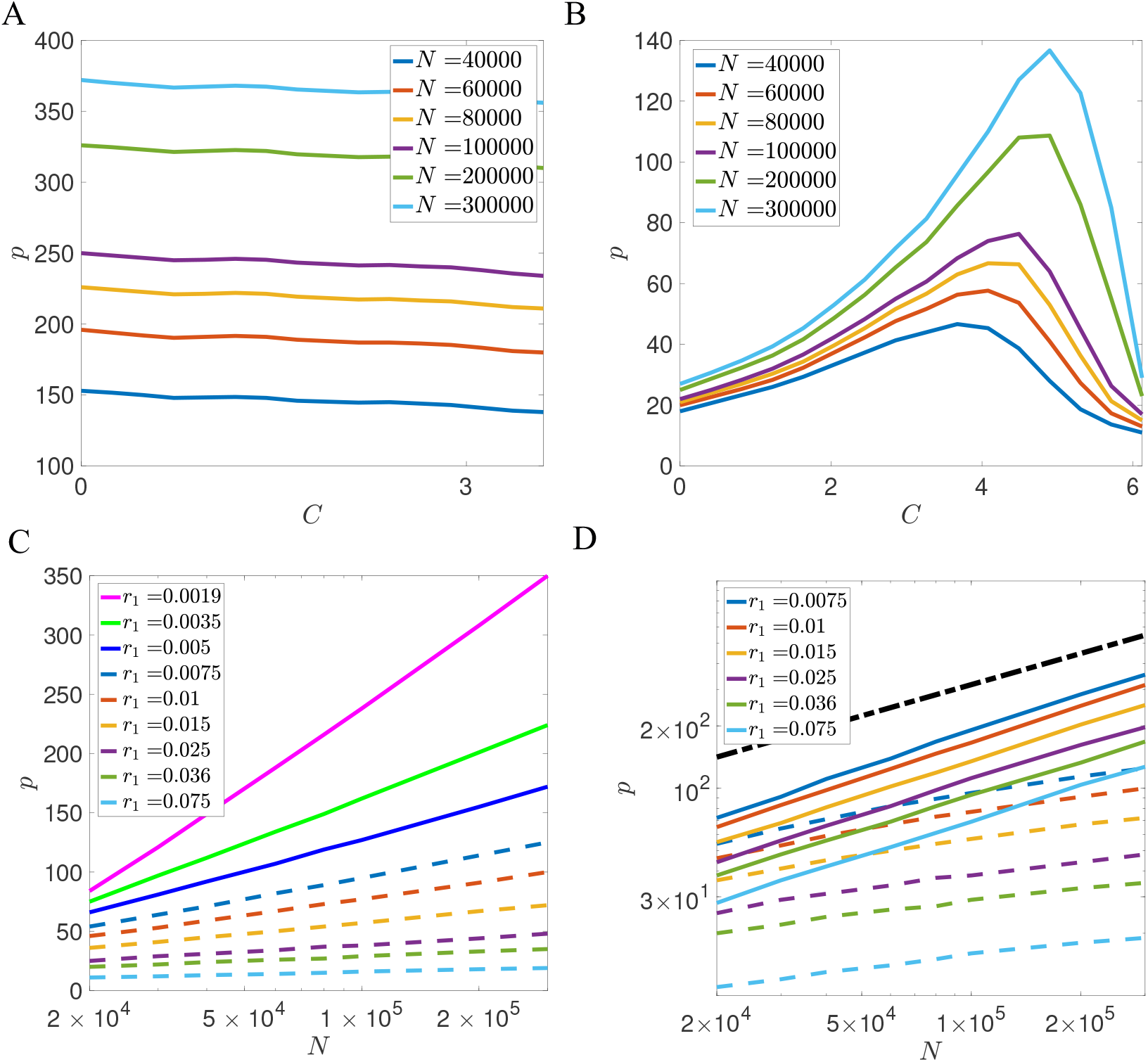
Dependence of storage capacity *p* on potential width *C* and network size *N*, for representative examples of decay rate *r*_1_. (A) Dependence of *p* on *C* for 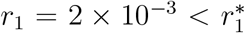. In the small *r*_1_ limit, the optimal potential width *C*^*^ is zero 0 (i.e. a single well potential), and *p* decreases weakly with *C*. The number of stored patterns increases logarithmically with network size, as demonstrated in (C). (B) Dependence of *p* on *C* for 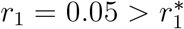. The number of stored patterns *p* reaches its maximum at a nonzero value of *C*. The optimal storage capacity increases as a power law of *N*, as demonstrated in (D). (C) Storage capacity of the single-well potential model (*C* = 0). Storage capacity *p* is plotted with network size *N* on a semi-log plot. The storage capacity decreases when *r*_1_ increases as indicated by Equation (10). Solid lines: examples when 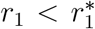. Dashed lines: examples when 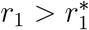. (D) Storage capacity of the double-well potential model (*C* = *C*^*^). Storage capacity *p* is plotted with network size *N* on a log-log plot. When 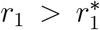, the optimal storage capacity is much larger than the storage capacity of the single well potential model (dashed lines with the same color). The dash-dotted black line represents 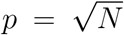 for reference. Other parameters in this figure are given as: *r*_2_ = 1, *r*_3_ = 0, *θ* = 0, *f* = 0.5, *c* = 0.05.

In the following, we will demonstrate that the storage capacity of the palimpsest model with fixed *τ* can be substantially enhanced by introducing the double-well potential in the synaptic dynamics.

#### Double Well Potential

The capacity of the model with double-well potential is influenced by the width of the potential wells *C*, and the model displays distinct behaviors depending on *r*_1_. When *r*_1_ is small, the storage capacity *p* is optimized for *C* = 0 (single well potential), but decreases only weakly with *C*, as shown in Figure 2A. In this case, the model is similar to the previously studied pure forgetting model [9], and the storage capacity only increases logarithmically with *N*, as demonstrated by the solid lines in Figure 2C.

However, when *r*_1_ is larger than a critical value 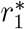, so that the potential significantly influences the dynamics of the weights in between presentations, the capacity is no longer maximized at *C* = 0. Rather, there is an optimal value *C*^*^ that maximizes capacity, as demonstrated in Figure 2B. When *C* = 0, the storage capacity increases logarithmically with the network size. However, the storage capacity at the optimal value of *C* increases much faster than the storage capacity with *C* = 0, as shown in Figure 2B. When *C* is increased beyond the optimal value, the storage capacity drops abruptly since jumps induced by inputs are too small to cross the boundary between the two wells. The optimal *C* depends on both *r*_1_ and *N*. We find that the critical value of *r*_1_ at which the optimal *C* becomes nonzero is 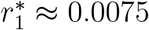 as shown in Figure 3A. In a broad parameter region, the optimal value *C* is larger than one, which means that multiple pattern presentations are required to induce the switch between two stable states of synapses. Our results also indicate that smaller *r*_1_ or larger *N* will lead to a higher value of the optimal *C*.

**Figure 3.**
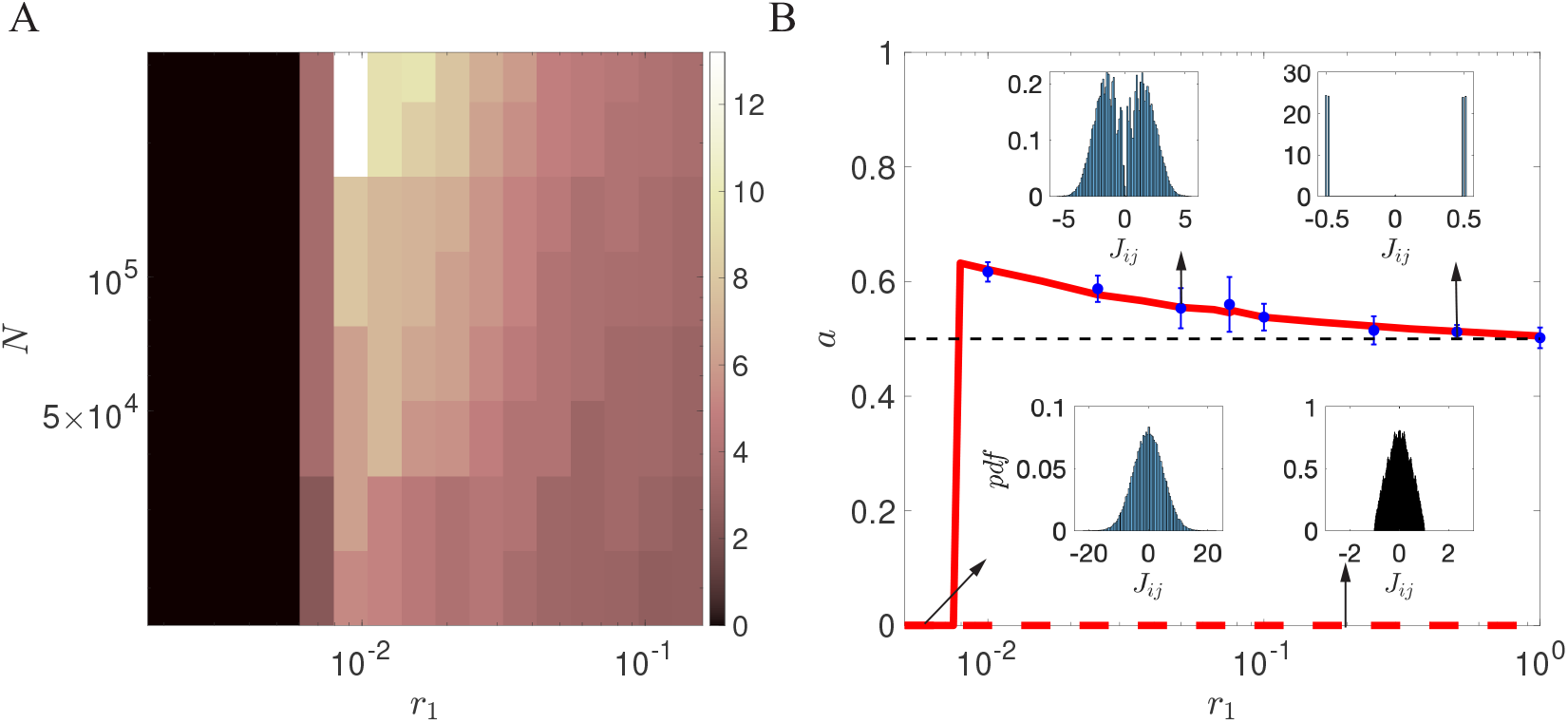
(A) Dependence of optimal potential width *C*^*^ on network size *N* and potential depth *r*_1_. There is a critical value 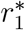 that separates two regimes: When 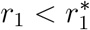, capacity is optimized for a single well potential, *C*^*^ = 0. When 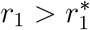, *C*^*^ is non-zero, and increases as *N* increases and *r*_1_ decreases, indicating more pattern presentations are required to induce the switch between two stable states of synapses. Our analytical and numerical computation suggest that 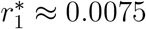, as indicated by the boundary of the black region. (B) Scaling exponent between *p* and *N* as a function of *r*_1_, with *r*_3_ = 0. Solid red lines represent: theoretical prediction for optimal *C*. Dashed red line: Theoretical prediction for *C* = 0. Blue circles: Simulation results for optimal *C* (mean and standard deviation computed over ten independent realizations). For 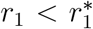, *p* ∼ log(*N*) gives *a* = 0. When 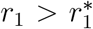, the model with *C* = *C*^*^ can has a power law dependence on *N* (red solid line), while the model with *C* = 0 gives *a* = 0 (dashed red line). The exponent *a* decreases with *r*_1_ and converges to 0.5 in the large *r*_1_ limit. Insets show distributions of synaptic weights for representative examples. The distribution of *J*_*ij*_s goes from unimodal when *C* = 0, to bimodal for the optimal *C* and 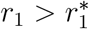 to quasi-discrete when *r*_1_ becomes large.

For each value of *r*_1_ and *N*, we focused on the value of *C* that maximizes capacity. For the region 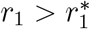, we found that the optimal storage capacity *p* with double-well potential can be well described by a power law of the network size *N*

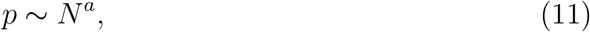

as indicated in Figure 2D. We show the dependence of the exponent *a* on *r*_1_ in Figure 3B, together with representative distributions of weights at a few values of *r*_1_. This plot shows three qualitatively different regions as a function of *r*_1_:

##### Shallow potential region

In the flat potential limit (i.e., 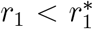), *p* has a logarithmical dependence with *N*, and thus the exponent *a* ∼ 0. The distribution of the weights in this region is unimodal. The model in this limit is equivalent to models with exponentially decaying memories [8, 40].

##### Intermediate region

When *r*_1_ increases beyond 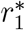, the storage capacity of the network can be significantly enhanced by optimizing the potential width *C*. In this region, the network exhibits a bimodal weight distribution, and the exponent *a* is slightly larger than 0.5 (solid line in Figure 3B), while the network with unimodal weight distribution (*C* = 0) still gives *a* = 0 (dashed line in Figure 3B).

##### Deep potential region

When *r*_1_ further increases, the weights are more and more attracted to the minima of the potential well in between pattern presentations, and the model becomes more and more similar to a binary synapse model. In the large *r*_1_ limit, the exponent *a* tends to 0.5, and the distribution of weights becomes close to the sum of two delta functions *δ*(*J* + *C*) + *δ*(*J* − *C*). This scenario is similar to the model with stochastic binary synapses studied in [7, 30].

### 3.2 Comparison between double well synapses and binary Markovian synapses

In models with discrete synapses, the transition between states is often described by a Markov process. Tsodyks [29] and Amit and Fusi [7, 30] studied the scaling of the number of stored patterns with network size *N* in networks with such synapses, and showed that the capacity scales as 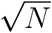 when transition probabilities are of order 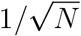. Fusi et al introduced a more general model with multiple states, and showed the stored information could decay more slowly with well-designed transition probability between states [32]. Lahiri and Ganguli derived an upper bound of the stored information for general Markov models [34]. We now turn to a comparison between double well models and such Markovian models. For a fair comparison between these two models, we binarize weights in the double well model,

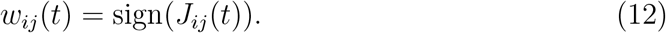

*w*_*ij*_(*t*) is thus a binarized connectivity matrix whose entries take +1 or −1 values. Similarly as in previous studies [41–43], we quantify information storage using a signal to noise ratio

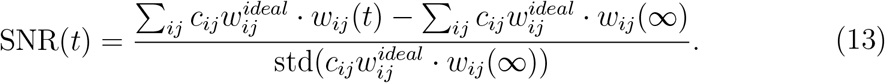

Here the standard deviation is calculated over all elements in the matrix, and the denominator is essential 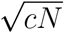 is the state of the connectivity matrix at *t* → ∞. 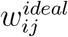 is the ideal state of the weight determined by the learning rule Equation (5). At time 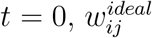 is 1 if it experiences a potentiation event and is −1 if it experiences a depression event (i.e., 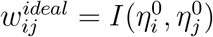 in Equation (5)).

For the general Markov model, SNR can be written as a sum of exponentials, as described by eq. (6) in [34]. The envelope of the SNR curve is given by eq. (19) in [34].

For the model with double well synapses, the transition probability between two states depends on the position of the weight within the potential well, which leads to non-Markovian effects. A weight that has been potentiated or depressed at a previous time without crossing the barrier is closer to the boundary between the two potential wells and is thus more likely to jump into a new state. In this model, the SNR can be calculated using (see Appendix III for details):

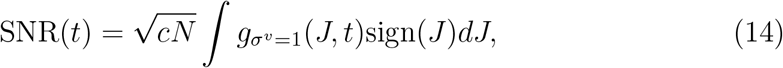

where *g*_+_(*J, t*) is the time-dependent distribution of the weights, conditioned on a potentiation event at time *t* = 0 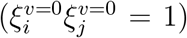, defined through eqs. (16) to (19) and (24).

In this analysis, we compare the SNR of the double-well model with that of a corresponding Markov model. The Markov model has two states represented by +1 and −1, and each weight has a probability of switching from −1 to +1 when it is potentiated (i.e., 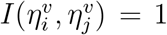), or from +1 to −1 when it is depressed (i.e., 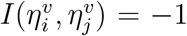). The transition probability between the two states is set to be the same as the transition probability in the double-well model. Therefore, the Markov model and the double-well model have the same initial SNR when a memory is learned, as illustrated in Figure 4.4. However, the SNR of the Markov model initially decays slowly and then rapidly drops off after *t* ∼ 10, while the SNR of the double-well model does not monotonically decrease due to the analog depth provided by the double-well synapses, which allows the synaptic state to have a rich dependence on neuronal activity history. Consequently, the SNR of the double-well model can surpass the upper bound of SNR in the Markov model and decay much more slowly than the SNR of the Markov model, resulting in an improved storage capacity, as shown in Figure 4A.

**Figure 4.**
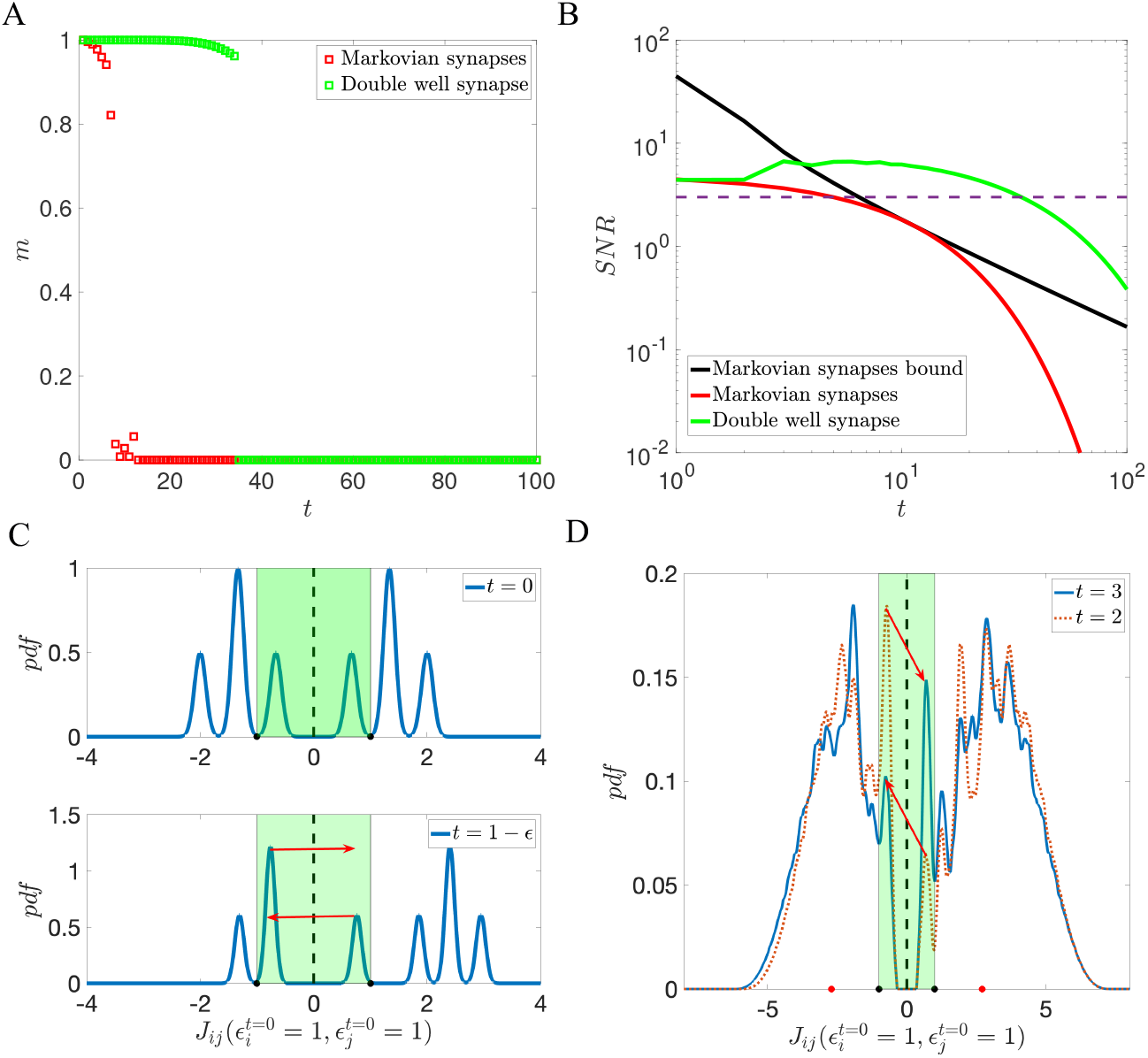
(A) Comparison between the storage capacity of Markovian binary synapses and binarized double well synapses defined in Equation (12). The parameters of the double well model are *r*_1_ = 0.1, *r*_2_ = 1, *r*_3_ = 0, *θ* = 0, *f* = 0.5, *cN* = 2000. The width of the potential *C* is chosen to maximize storage capacity. The Markov model is chosen with the same transition probability as the double-well model. (B) Comparison between SNR of Markovian synapses and binarized double well synapses, as a function of time elapsed since the presentation of a given pattern. Green curve: SNR of the double well model calculated using eq. (14). Red line: SNR of the Markov model with the same transition probability. Black line: Upper bound of SNR for two-state Markov models. The SNR and the upper bound of SNR for the Markov model are calculated using eq. (3) and eq. (19) in [34]. Dashed line: critical SNR below which the memories are no longer retrievable. The intersection between the SNR curve and this dashed line determines the storage capacity of the network.(C) Cartoon of temporal evolution of distribution of synapses that undergo potentiation at *t* = 0, i.e. those for which 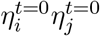) = 1 just before potentiation (top) and after (bottom). The highlighted region denotes the region around the maximum of the potential where synapses that reside in one of the wells can make a transition to the other well (*r*_2_ = 1). Just before potentiation (*t* = 0), the distribution is symmetric. The distribution is then shifted up by *r*_2_, before decaying towards the respective wells. As the distribution becomes asymmetric near 0 (within the range of 0 *± r*_2_), the next presented uncorrelated patterns will cause more probability mass to shift towards the positive side than towards the negative side, as demonstrated in the bottom figure. (D) Distribution of connections that have been potentiated at time *t* = 0, at later times *t* = 2 and *t* = 3. Black dots indicate *±r*_2_ = *±*1, while red dots indicate the minima of the potential, *±C* = *±*2.7. As demonstrated in (C), an uncorrelated pattern leads to a larger probability of switching from the low well to the high well than the opposite transition, resulting in an increase in SNR from *t* = 2 to *t* = 3, as seen in (B). At longer times, the distribution eventually returns to a symmetric distribution, leading to a decrease in SNR.

To better understand the history-dependent nature of double-well synapses, we show the changes in the distribution of potentiated connection 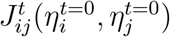 over time in Figure 4C. After a pattern is learned, the distribution of 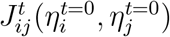 becomes asymmetric, and subsequent uncorrelated patterns can cause a probability flux towards the positive side, as demonstrated in Figure 4C. This happens in particular for the parameters of Figure 4 at *t* = 2, which leads to an increased SNR from *t* = 2 to *t* = 3, as seen in Figure 4B. After a sufficient amount of time, the distribution eventually returns to a symmetric asymptotic distribution, resulting in a decay of the SNR towards zero.

### 3.3 Robustness of stored information to noise

Neural circuits are required to not only learn new information but also to retain old memories. Recent experiments suggest that a significant fraction of synaptic plasticity is random noise, independent of learning [44, 45]. How can neural networks maintain long-lasting memories in the presence of noise? One long-lasting hypothesis is that discrete-like synapses could benefit neural networks by increasing their robustness with respect to noise (see e.g. [46]). Here, we can verify this hypothesis in the model with double-well synapses by adding noise to the model. This noise is described by the last term in Equation (3), where *r*_3_ quantifies the magnitude of the noise term.

We first investigated how the potential width affects the network’s storage capacity in the presence of noise. We set the potential depth to a fixed value of *r*_1_ = 0.1 and calculated the storage capacity for every combination of *r*_3_ and *C*. As illustrated in Figure 2B, there is an optimal value *C*^*^ that maximizes the storage capacity of the network without noise. If the potential width increases beyond *C*^*^, the Hebbian learning term defined in Equation (5) is insufficient to induce the transition between different states, resulting in an abrupt decrease in storage capacity. For models with *C < C*^*^, the network’s storage capacity decreases monotonically with the noise strength *r*_3_, as shown in Figure 5. However, for models with *C > C*^*^ (e.g., *C* = 4, *C* = 5), there is a peak in the storage capacity as *r*_3_ increases, indicating that noise facilitates the learning of new memories.

**Figure 5.**
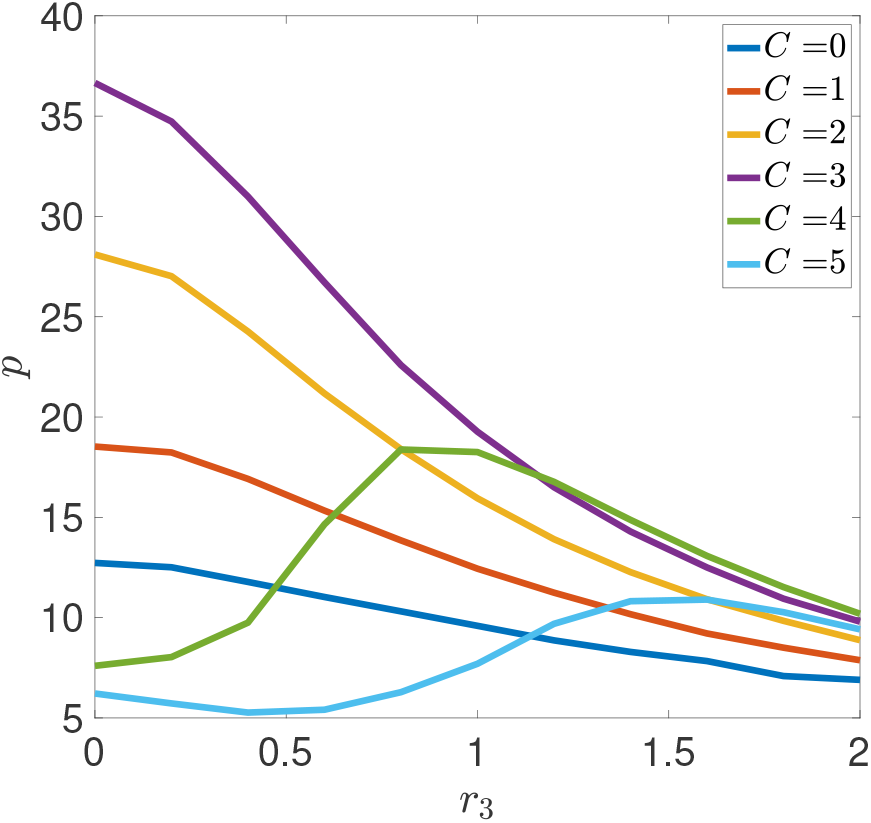
Dependence of Storage Capacity on Noise Strength *r*_3_ for different values of Potential Width *C*. When *C* is small (e.g., *C* = 0, 1, 2, 3), the storage capacity *p* monotonically decreases with the noise strength *r*_3_. However, for larger values of *C* (e.g., *C* = 4, 5), the storage capacity first increases with noise intensity, reaches a peak and then decreases. Thus, noise facilitates the learning of new memories in such cases. Other parameters are given as *N* = 10, 000, *r*_1_ = 0.1, *r*_2_ = 1, *θ* = 0, *f* = 0.5, *c* = 0.05.

When *C* is large, the Hebbian learning term is insufficient to induce transitions of sufficient numbers of synapses, leading to an inability of the network to learn new memories. Our findings suggest that noise can help with the learning of new memories in such situations. This scenario is comparable to the model presented in [7, 30]. In this way, neural circuits can learn memories without requiring extreme precision in synaptic dynamics.

We also analyzed the effect of potential depth on the robustness of stored memories. For each fixed potential depth *r*_1_, we can choose the potential width *C* that optimizes the storage capacity. With the optimal *C*, the storage capacity of the network always decreases when the noise strength increases, as shown in Figure 6. However, the storage capacity decreases more slowly if the potential is deeper, which indicates the model is more robust to input noise. To quantify this effect, we introduced a robustness parameter as

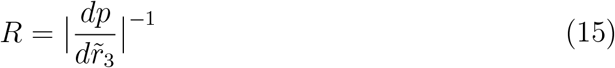

where 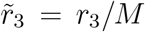 and *M* is the standard deviation of weight strength defined in eq. (23). 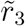 measures the perturbation to the weight caused by the input noise. A larger *R* means the storage capacity decreases more slowly in the presence of input noise.

**Figure 6.**
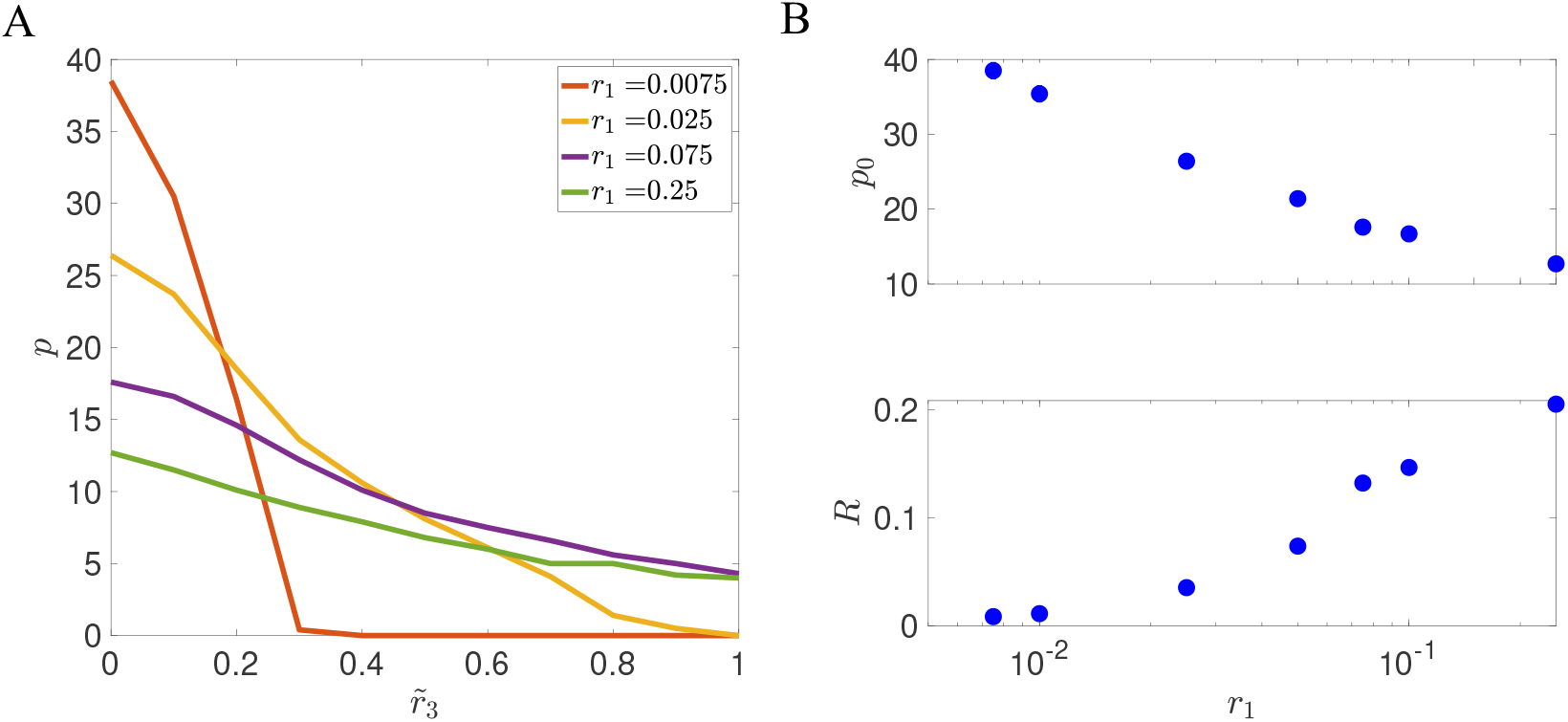
Trade-off between capacity and robustness when varying potential depth *r*_1_. (A) The storage capacity decrease when the noise strength increases strongly depends on *r*_1_. Small values of *r*_1_ lead to large storage capacity, but storage is highly sensitive to noise. Large values of *r*_1_ leads to small capacity, but storage is highly robust. (B) Trade-off between *p*_0_ and *R*, where *p*_0_ is the storage capacity in the absence of noise (*r*_3_ = 0), and *R* quantifies robustness to noise, Equation (15). With deeper potentials, the storage capacity is smaller but storage is more robust. Here, *C* is optimized for each value of *r*_1_ and other parameters are given as: *N* = 10, 000, *r*_2_ = 1, *θ* = 0, *f* = 0.5, *c* = 0.05.

We explored how the robustness *R* depends on the depth of the potential *r*_1_. As shown in Figure 6, when *r*_1_ increases, the storage capacity of the network in the absence of noise decreases. However, the robustness *R* becomes larger, indicating a trade-off between storage capacity and robustness as potential wells become deeper. From a biological perspective, each minimum of the potential can correspond to a cluster of receptors [5]. The parameter *r*_1_ may be related to the rate at which these clusters form. The timescale *τ* = 1*/*2*r*_1_ provides synapses with the analog depth to retain the history information of neuronal activity. Our results indicate that the balance between the memory storage capacity and the robustness of stored information would lead to an optimal *r*_1_.

### 3.4 Learning dynamics in the sparse coding limit

We next turn to a more biologically realistic scenario in which neurons are described by 0,1 variables, and the probability that a neuron is active in a given pattern, *f*, is *f* ≪ 0.5. We study the storage capacity of the network with double-well synapses with the learning rules described in Equations (6) and (7) in the sparse coding limit *f* ∼ ln *N/N* [47]. In this limit, it was shown that networks with binary synapses and on-line learning have a storage capacity that is comparable to the optimal capacity, unlike with other scalings of *f* with *N* where the capacity is sub-optimal [7, 47–49]. We first focus on the noiseless case (*r*_3_ = 0). For each value of *r*_1_, we optimize the storage capacity by adjusting the neuronal threshold *θ* and the potential width *C*. The relationship between the storage capacity *p* and the network size *N* is shown in Figure 7 for different choices of the learning rule and different potential depths *r*_1_.

**Figure 7.**
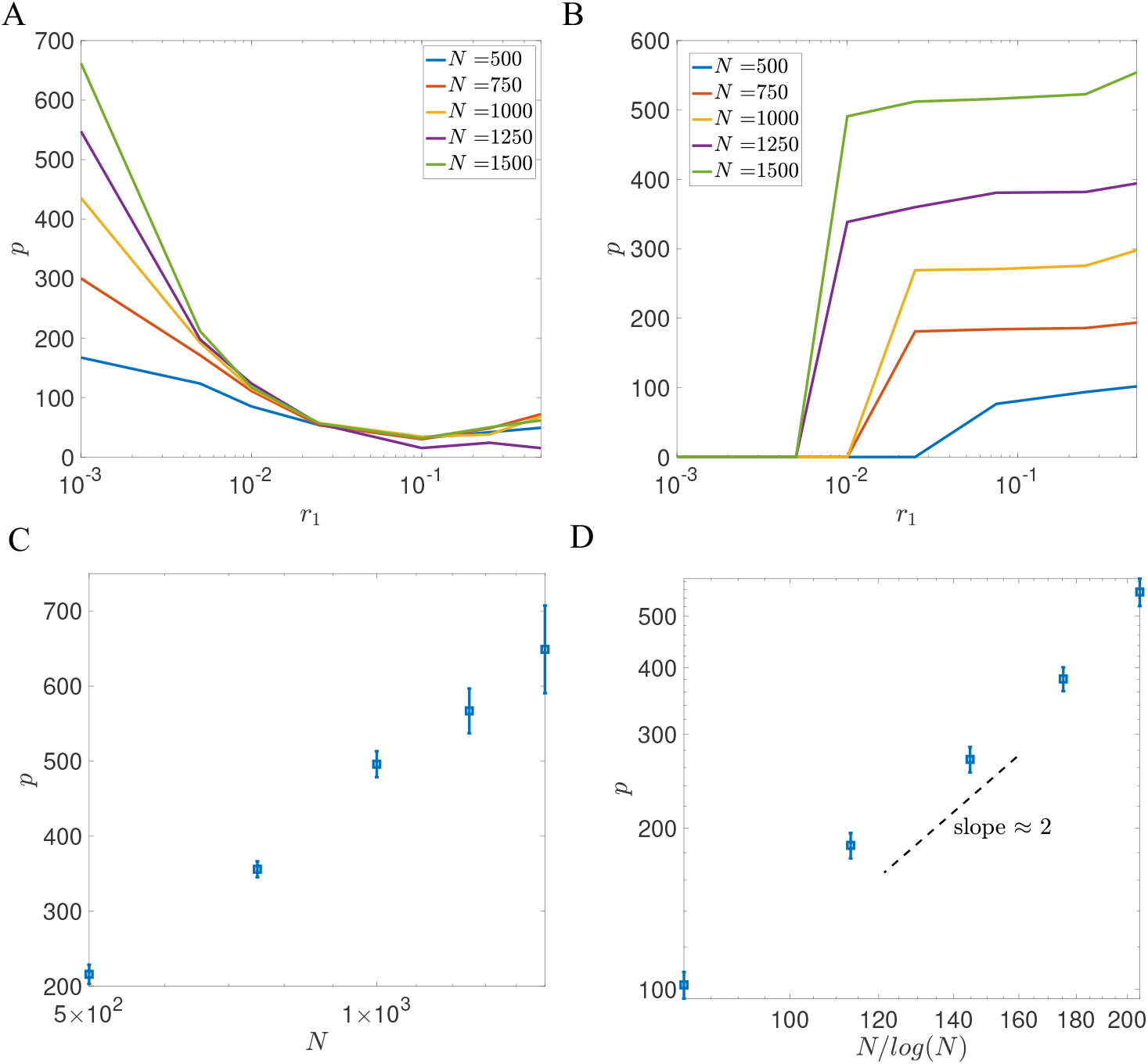
Storage capacity of the model with double–well synapses in the sparse coding limit. (A)(B) shows the storage capacity of the network with the learning rule defined in Equation (6) (A) and the learning rule defined in Equation (7) (B). (C) shows the dependence between *N* and *p* in the flat potential case (*r*_1_ = 10^−3^) for model with learning rule defined in Equation (6). Optimal *p* is plotted as a function of *N* in a semi-log plot. (D) shows the dependence between *N* and *p* in the deep potential case (*r*_1_ = 0.5) for model with the learning rule defined in Equation (7). In all curves, each data point is averaged over 10 realizations in all panels, and the error bar represents the standard deviation. *C* and *θ* are chosen to optimize for storage capacity. Other parameter are given as: *r*_2_ = 1, *r*_3_ = 0, *c* = 1, *f* = 4 log(*N*)*/N*.

We start with the learning rule defined in Equation (6), which is similar to the Tsodyks-Feigel’man learning rule [10, 36]. For this learning rule, we found that the optimal *C* is zero for all *r*_1_, and the distribution of synaptic strengths is unimodal. As demonstrated in Figure 7A-C, the storage capacity of the network increases logarith-mically with the network size *N*, with a prefactor that is proportional to *τ* = 1*/*(2*r*_1_) as indicated by Equation (10). This case is similar to the model with continuous synapses and exponentially decaying memories, as discussed in [8, 9, 37].

In contrast, the learning rule described in Equation (7), which is similar to the rule introduced by Amit and Fusi [7], exhibits a large storage capacity when the potential is deep, as illustrated in Figure 7B. In the deep potential regime, the weight distribution becomes strongly bimodal, with two peaks centered around *±C*.

When there is a coincidence of pre and post-synaptic activity, the learning rule in Equation (7) allows the synaptic connection to cross the potential barrier with probability ∼1. When one neuron is active while the other is silent, synapses switch to the low state only with low probability. This scenario is similar to the model discussed in [7]. As shown in Figure 7D, the storage capacity of this learning rule leads to *p* ∼ (*N/* log(*N*))^2^ in the sparse coding limit.

Our result demonstrates that the double-well model can achieve a much larger storage capacity in the sparse coding scenario than when memories are densely coded, similarly to previously studied models. In the flat potential case, the storage capacity scales as *p* ∼ log(*N*), but with a prefactor that increases when coding becomes sparse. However, in the deep potential case, the capacity scales supralinearly with network size, *p* ∼ (*N/* log(*N*))^2^, in the sparse coding limit, while it is sublinear when coding is dense.

We also investigated the robustness of the network with learning rules defined in Equations (6) and (7) in the sparse coding case. We explored how the storage capacity changes in the presence of noise, in both shallow and deep potential scenarios. As shown in Figure 8, when the noise strength *r*_3_ increases, the storage capacity of the network with deep potentials decays much more slowly than the storage capacity of the network in the flat potential case. This result is consistent with the result for the balanced input case. Notice that for the given network size *N* = 1, 500, the storage capacity in the deep potential case is smaller than the storage capacity in the flat potential case for small *r*_3_, despite the fact that it should increase faster with *N*. This is because the flat potential case has a much larger decay constant, *τ* = 1*/*2*r*_1_, leading to a larger prefactor in the relationship between *N* and *p* as indicated in Equation (10).

**Figure 8.**
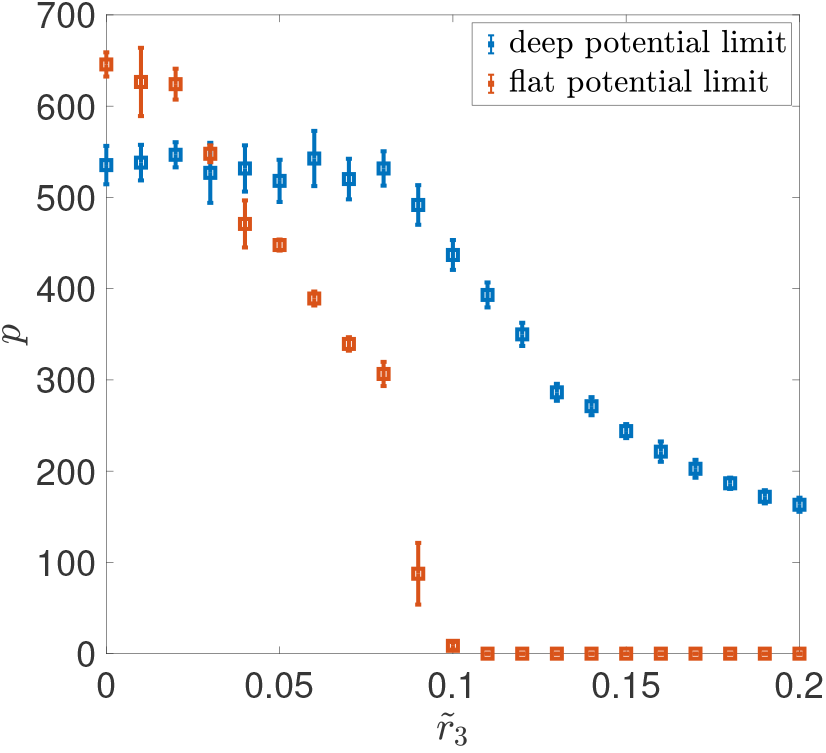
Robustness of the double well model in the sparse coding case. Orange squares represent the storage capacity for the model with the learning rule defined in Equation (6) in the flat potential case (*r*_1_ = 10^−3^). Blue squares represent the storage capacity for the model with the learning rule defined in Equation (7) in the deep potential case (*r*_1_ = 0.5). Each point is averaged over 10 independent realizations, and the error bar represents the standard deviation. Other parameters are given as: *r*_2_ = 1, *c* = 1, *N* = 1, 500, *f* = 4 log(*N*)*/N*.

## 4 Discussion

A long-standing question in neuroscience is whether individual synaptic plasticity events are better described as small changes on a continuum of possible values that are then stable over long time scales, or as large changes among a small discrete set of states [6, 26, 50]. These two scenarios have been studied essentially independently of one another, in networks with unconstrained analog synapses [1,8,36], or in networks with binary synapses [7, 29–31].

In this study, we introduced a model that can interpolate between these two scenarios. Synapses in our model evolve in a double well potential, whose shape is described by its depth *r*_1_ and width *C*. When the depth goes to zero, the model becomes equivalent to classic models with continuous synapses, while in the opposite limit of a large depth, the model becomes similar to models with binary synapses. In the absence of inputs, a synapse decays to one of the potential minima exponentially, with a time constant *τ* = 1*/*2*r*_1_. External input can cause synapses to jump from one well to the other, leading to a stable change in the absence of noise or additional external inputs leading to subsequent jumps. When the size of the jumps is smaller than the distance between the minimum of a well and the barrier that separates the two wells, repetitive stimulation is needed for a synapse to jump from one well to the other. The number of required stimuli is determined by the potential width *C*. This dynamics is reminiscent of phenomena such as synaptic consolidation or late long-term potentiation. Experimental studies on plasticity in the hippocampus have shown that weak stimulation leads to synaptic changes that decay to their initial value in a relatively short period. However, a strong stimulation results in an enhanced synaptic connection, that persists for the entirety of the recording [51]. This synaptic consolidation process involves multiple complex biochemical dynamics, and our double-well potential model could be seen as a minimal implementation of this idea in the framework of attractor neural networks.

There are multiple lines of evidence that suggest that synapses have a small set of stable states. Synaptic plasticity has been shown to be implemented by the addition of unitary synaptic ‘nanomodules’ to dendritic spines, the loci of excitatory synapses onto pyramidal cells, the main excitatory neuronal type in the cortex and hippocampus [5]. These nanomodules contain spatially localized clusters of pre- and postsynaptic proteins that are aligned with each others. These clusters form due to a reversible diffusion-trapping process of receptors and scaffold proteins on the membrane [52]. Our model can be seen as a minimal model for this scenario, where the two minima of our potential well could correspond to spine configurations with one and two clusters of receptors, which represent close to 90% of the observed spines in [5]. The depth of the potential could be related to the rate at which these clusters form, which depends on various factors, such as the fluidity of the synaptic membrane and the affinity of the receptors for scaffold proteins. The potential width *C* determines the synaptic plasticity gap, describing the number of stimuli required to induce the formation of new nanomodules. The idea of discrete stable states is also consistent with electron microscopy data showing distributions of spine sizes in cortex are bimodal [6], and models of synaptic plasticity relying on biochemical interaction networks of the post-synaptic density [53–55].

Our results show that a network in which synapses have two potential wells, and consequently there is a bimodal distribution of synaptic strengths, has a larger storage capacity than a model in which synapses evolve in a single well, and therefore the distribution of weights is unimodal, provided 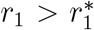 or equivalently 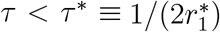. Furthermore, our results show that *C*^*^, the optimal potential width, is greater than one whenever *r*_1_ is larger than its critical value, as shown in Figure 3A. The optimal value of *C* increases with both network size *N* and the time constant *τ*. For large networks, this implies that many repetitive stimuli are needed to induce transitions between potential wells. In this regime, the storage capacity of the network increases as a power law of the network size, instead of a logarithmic relationship. The exponent is close to 0.5 for densely encoded memories, but it becomes equal to 2 (with logarithmic corrections) with sparsely encoded memories when the number of active neurons per memory is logarithmic in network size. These exponents are similar to networks with binary synapses [7, 31]. Our results also suggests that synaptic noise can facilitate the learning of new memories, in situations in which that the Hebbian learning term is insufficient to induce transitions between potential wells. Overall, this suggests that a network with such synapses can store memories robustly without requiring high precision in synaptic dynamics.

Our model captures the process of synaptic consolidation in an abstract fashion, as the decay of the synaptic state towards one out of two potential minima. The decay is characterized by a single time constant, *τ*, which results in a storage capacity that depends sublinearly on the network size in the balanced input case. Experimental and theoretical work have revealed that synaptic plasticity contains several phases across multiple time scales [56–60]. It will be interesting to generalize the model we have proposed to include dynamics with multiple timescales, which would account for various biochemical processes during synaptic consolidation. Such an extension may increase the storage capacity of the network, leading to an almost linear dependence on network size, as suggested by previous research [61]. The phenomenon of synaptic consolidation can be well described by the theory of synaptic tagging and capture [56,58,62]. According to this theory, stimulation can result in the tagging of synapses. When weak stimulation is applied, the tag gradually decays. Strong stimulation instead can tag more synapses, initiating the production of plasticity-related proteins that lead to the consolidation of the tagged synapses. Although our double-well synapse model has some similarities with the theory of synaptic tagging and capture, it does not account for a number of biological details. For instance, our model only considered independent synapses, while in reality, different synapses on the same dendritic tree can share tagging, since plasticity-related proteins are synthesized in the soma of postsynaptic neurons, which are available to all presynaptic pathways. Moreover, experiments suggest that synapses have varying stimulation thresholds for discrete changes in their efficacy [26], while we assumed homogeneous synapses. Future research will be necessary to include these biological details or investigate these phenomena with more realistic neuron and network models. We also note that other forms of memory consolidation are likely to play an important role. A recent study demonstrated that adding a pattern reactivation mechanism in attractor neural networks with synapses evolving in a single well potential, mimicking systems memory consolidation [63], can also produce power law scaling storage capacity and lead to lifelong learning scenarios [9]. It will be interesting to incorporate synaptic consolidation and pattern reactivation together in an attractor neural network to explore how double-well synapses affect memory function in the context of lifelong learning.

## Appendix

### I. The probability density function of synaptic weights

The goal of this section is to derive an equation for *g*(*J, t*), the probability density of synaptic strength *J* at time *t*, and distributions of synaptic strengths conditioned by inputs experienced in the past. From Equation (3), one can derive a master equation for *g*(*J, t*):

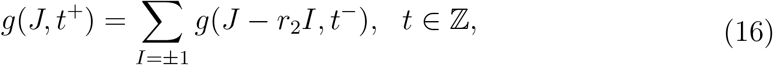

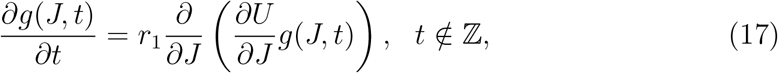

where *t*^−,+^ denote times right before or after a new pattern is presented. With the piece-wise parabolic potential defined by Equation (4), Equation (17) yields

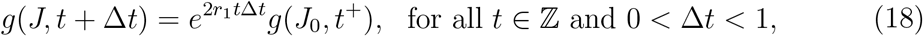

where

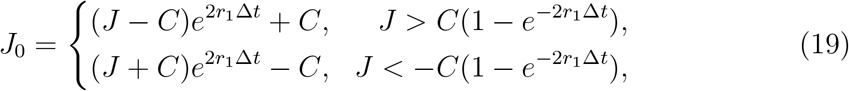

After a sufficiently long time, the distribution of synaptic weights should only depend on the time elapsed since the last input presentation, and should therefore be periodic with period 1:

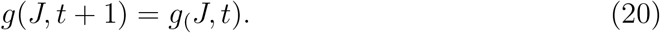

In addition, the probability density function should satisfy the usual boundary conditions,

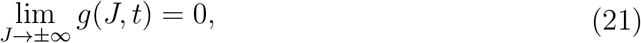

and the normalization condition

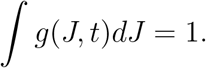

Using eqs. (16) to (21), one can solve the asymptotic distribution of the weights right before a new pattern is presented, *g*_∞_(*J*) ≡ *g*(*J, t*^−^) for sufficiently large *t* ∈ Z → ∞. The mean and root mean square (RMS) of asymptotic distribution *g*_∞_(*J*) are given as

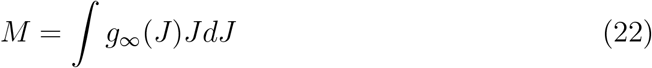

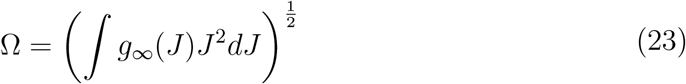

We next turn to the distribution of synaptic strengths, conditioned on plasticity events at time *t* = *v* and *t* = *u*. We use the *η*^*u*^ to denote the most recently presented pattern and *η*^*v*^ to denote the previously stored pattern where *v < u*. Distributions of synaptic strengths at time *t*, conditioned on a plasticity event at *t* = *v*, are denoted by 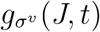, where *σ*^v^ = *I*(*v*) = *±*1 denotes potentiation or depression. Just after the presentation of the pattern, these distributions are given by:

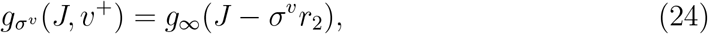

The distributions then evolve according to eqs. (16) and (17), replacing *g*(*J, t*) by 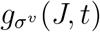. At any given time, one can calculate the mean 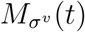 and RMS 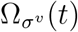 of the weight distributions 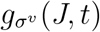:

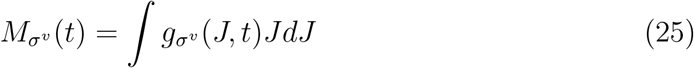

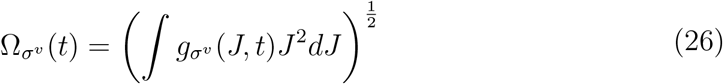

Distributions of synaptic strengths at the time *t* = *u*, conditioned plasticity events at time *t* = *v* and *t* = *u* are denoted by 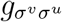, where *σ*^*v*^ = *I*(*v*) = *±* 1 and *σ*^*v*^ *I*(*u*) = *±*1. The distribution of 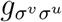 is given

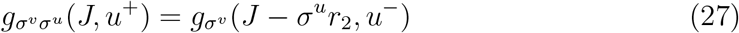

by Its mean and RMS are

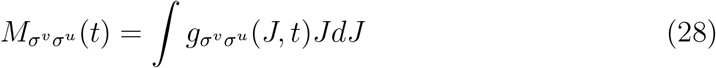

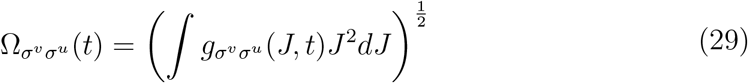

### II. Storage capacity of the double well potential model

We now turn to the question whether at time *u*, the network is still able to retrieve a pattern *η*^*v*^ that was stored at a previous time *v < u*. Here, we focus on the standard coding (‘balanced input’) scenario *f* = 0.5, and *θ* = 0. Simulations show that sometimes, when initialized at a pattern stored in the past, the network mistakenly retrieves the most recently shown pattern *η*^*u*^ instead [37]. To calculate the storage capacity, we thus need to define two order parameters:

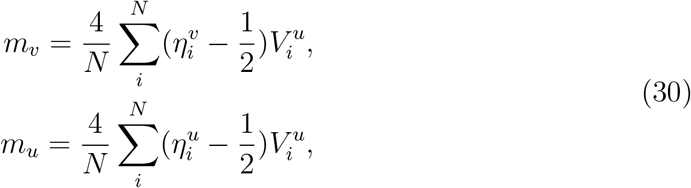

where 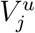 represents the fixed point of eq. (1) with initial condition 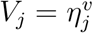, *u > v* ∈ ℤ and *η*^*u*^ represent the most recently shown pattern.

When the network has overlaps *m*_*v*_ and *m*_*u*_ with patterns shown at times *u* and *v*, the probability that a neuron is in a given state *V*_*i*_, conditioned on 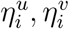 can be written as:

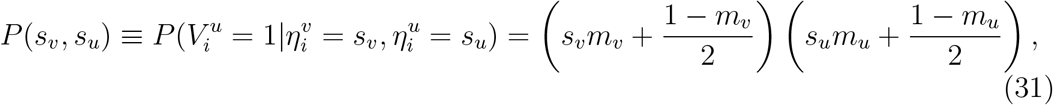

where *s*_*v*_, *s*_*u*_ ∈ {0, 1}. To proceed, we need to compute the distribution of local fields conditioned on the state of the neuron in patterns shown at times *u* and *v*. The local fields are given by

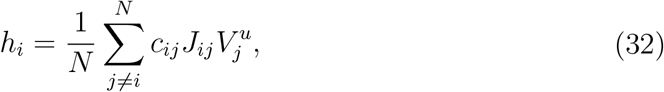

where in the right hand side, the distribution of *J*_*ij*_ are given by 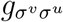 where 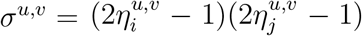, and the distribution of 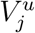 are given by eq. (31). For large *cN*, (*cN* ≫ 1), we can use the central limit theorem and approximate the local field distributions by a normal distribution 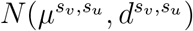, where 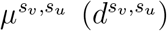 is the mean(variance) of the local field 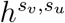, conditioned on 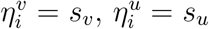. These two moments are given by

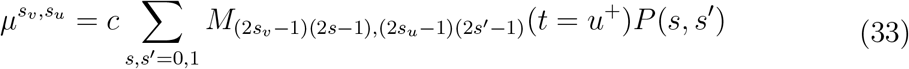

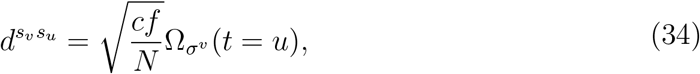

where *s*_*v*_, *s*_*u*_, *s, s*^*′*^ ∈ *{*0, 1*}*. The order parameters *m*_*v*_, *m*_*u*_ can be written as:

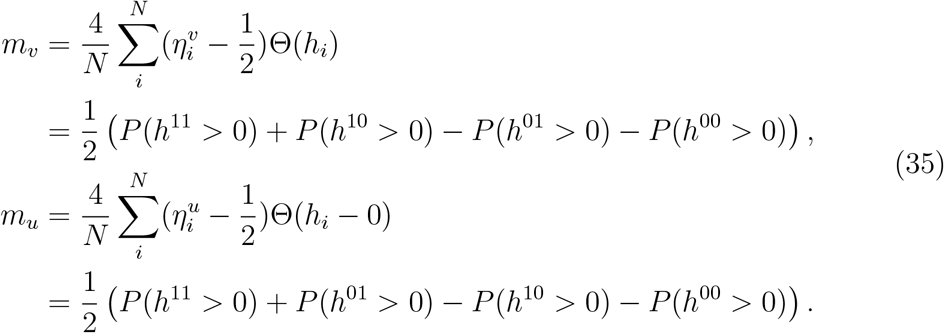

In the balanced input case, 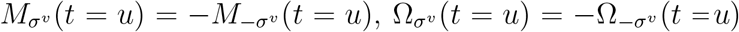 and thus

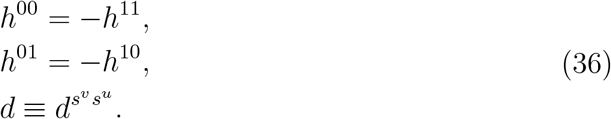

The relation Equation (36) can be simplified to:

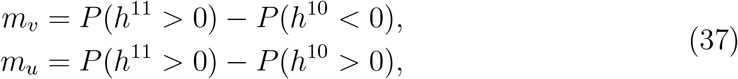

Use the notation defined in eqs. (33) and (34), we obtain self-consistent equations for *m*_*v*_ and *m*_*u*_:

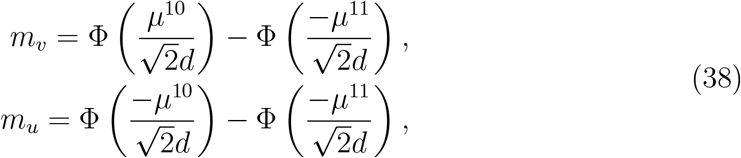

where 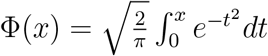. eq. (38) can be solved numerically by updating eqs. (31), (33) and (34) using *m*_*v*_, *m*_*u*_. the pattern *η*_*v*_ is retrievable if there exists a solution *m*_*v*_ ∼ 1, *m*_*u*_ = 0, that is stable to small perturbations *m*_*u*_ = *ϵ*. In order to solve this mean-field equation, we convert eq. (38) to a map, in which the values of *m*_*u,v*_ at a given time step are given by the r.h.s. 0f eq. (38) where *μ*^11,10^ and *d* are computed using the values of *m*_*u,v*_ at the previous time step. To check the stability of the state with *m*_*v*_ *>* 0, *m*_*u*_ = 0, we take as initial conditions *m*_*v*_ as 1 and *m*_*μ*_ as *ϵ*. We then proceed to update the values of *m*_*v*_ and *m*_*μ*_ iteratively until convergence is reached. In practice, we used *ϵ* = 5 *×* 10^−2^.

### III. Signal-to-Noise Ratio

We focus on the calculation of the signal-to-noise ratio in the balanced input case. In this case, *w*_*ij*_ has the equal probability to be +1 and −1, thus we have:

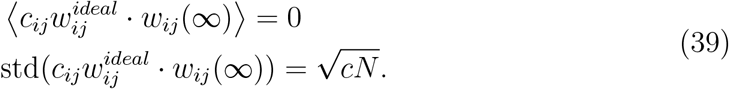

Therefore, the SNR can be simplified as follows:

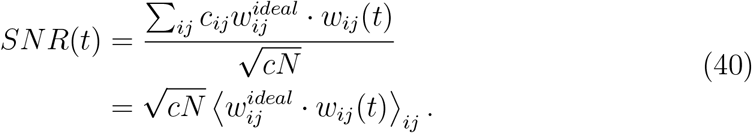

*w*_*ij*_(*t*) is drawn from the distribution of 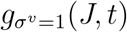 and the distribution of 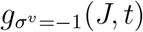 with the same probability in the balanced input case, then the average 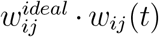 can be calculated by:

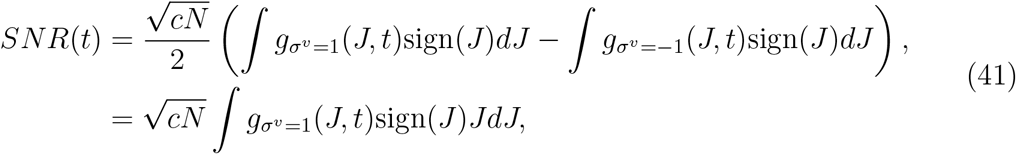

where 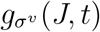 are defined through eqs. (16) to (19) and (24), and we used the identity 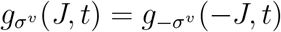 in the balanced input case.

